# Differential alteration of plant functions by homologous fungal candidate effectors

**DOI:** 10.1101/2020.10.30.363010

**Authors:** Karen Cristine Goncalves dos Santos, Gervais Pelletier, Armand Séguin, François Guillemette, Jeffrey Hawkes, Isabel Desgagné-Penix, Hugo Germain

## Abstract

Rust fungi are plant pathogens that cause epidemics that threaten the production of important plant species, such as wheat, soy, coffee and poplar. *Melampsora larici-populina* (*Mlp*) causes the poplar rust and encodes at least 1 184 candidate effectors (CEs), however their functions are poorly known. In this study, we used *Arabidopsis* plants constitutively expressing CEs of *Mlp* to discover processes targeted by these fungal proteins. For this purpose, we sequenced the transcriptome and used mass spectrometry to analyse the metabolome of *Arabidopsis* plants expressing individually one of the 14 selected CEs and of a control line. We found 2 299 deregulated genes across the experiment. Among the down-regulated genes, the KEGG pathways “MAPK signaling pathway” and “Plant-pathogen interaction” were respectively over-represented in six and five of the 14 transgenic lines. Moreover, genes related to hormone response and defense were down-regulated across all transgenic lines are. We further observed that there were 680 metabolites deregulated in at least one CE-expressing transgenic line, with highly unsaturated and phenolic compounds enriched in up-regulated metabolites and peptides enriched among down-regulated metabolites. Interestingly, we found that transgenic lines expressing unrelated CEs had correlated patterns of gene and metabolite deregulation, while expression of CEs belonging to the same family deregulated different genes and metabolites. Taken together, our results indicate that the sequence of effectors and their belonging to families may not be a good predictor of their impact on the plant.

**Importance:** Rust fungi are plant pathogens that threaten the production of important crops, including wheat, soy, coffee and poplar. Effectors are used by pathogens to control the host, however in the case of *Melampsora larici-populina*, the causal agent of the poplar rust, and other rust fungi these proteins are poorly known. We used *Arabidopsis* plants expressing candidate effectors (CEs) of *Mlp* to better understand the interaction between this pathogen and its hosts. We found that expression of unrelated CEs led to similar patterns of gene and metabolite deregulation, while transgenic lines expressing CEs belonging to the same family showed different groups of different genes and metabolites deregulated. Thus, our results suggest that functional annotation of effectors based on sequence similarity may be misleading.

## Introduction

Plants have to defend themselves against different types of pathogens. Their first line of defense consists of passive barriers, such as the cuticle and cell wall, which prevent pathogens from entering the plant tissue and its cells. Upon successful entry of a pathogen, conserved pathogenic motifs, called Microbe-Associated Molecular Patterns (MAMPs), may be detected and activate the Pattern-Triggered Immunity (PTI) [1]. PTI includes the transient accumulation of reactive oxygen species (ROS), callose deposition, alteration of hormone networks and activation of defense genes [2, 3]. Finally, microorganisms secrete effectors into their host to modulate the host metabolism in favor of the pathogen. If detected, these effectors will activate the Effector-Triggered Immunity (ETI), leading to plant cell death in order to avoid pathogen spreading to surrounding cells [4].

Rust fungi are the largest group of fungal plant pathogens, infecting ferns, gymnosperms and angiosperms and causing important losses in food production [5, 6]. They are obligate biotrophs, produce two to five types of spores and infect one or two unrelated species to complete their life cycle [6]. To guard themselves against the defense mechanism of two different host species and to be able to feed on them, rust fungi deploy a large arsenal of effectors. To better comprehend the interaction between these pathogens and their hosts, and to provide new mechanisms to target in order to improve plant immunity, it is imperative that we understand how these effectors are secreted into host cells, how they evolve and how they act to promote pathogen growth [7, 8]. While the precise number of *bona fide* effectors carried by each rust fungi species is unknown, Duplessis and colleagues [9] established that the poplar rust (*Melampsora larici-populina*) genome encodes 1 184 small secreted proteins (SSPs) whereas the wheat stripe rust (*Puccinia graminis* f. sp. *tritici*) genome encodes 1 106 SSPs [9], which are considered candidate effectors (CEs). These CEs are grouped within families based on sequence homologies [10, 11]. Furthermore, effectors in the same family have been shown to interact with homologous R-proteins [12], however the virulence function of these effectors has seldom been investigated.

Previous studies have proposed different criteria to screen the genome of plant pathogenic fungi for high-priority CEs, including having less than 300 amino acids, high cysteine content, being expressed in infection structures during host infection or being detected in the host tissue during infection [11, 13, 14]. Once identified, putative effectors must be functionally characterized. In pathogens that are not obligate biotrophs, this can be achieved by silencing or overexpressing the gene encoding the CEs and analysing the outcome of an infection [15, 16]. For rust fungi and other obligate biotrophs, which are not amenable to genetic transformation, this direct investigative approach is not possible. The alternative solution proposed by different research groups is to use heterologous systems, either by transforming model plants to express the CE-encoding gene or by infecting model plants with pathogens able to express these genes [17, 18]. This way, it is possible to evaluate if immunity is compromised, as it was shown that effectors expressed in heterologous systems conserve their capacity to alter the plant’s susceptibility to pathogens [19-24]. The stable and transient expression of CEs from *M. larici-populina* in *Arabidopsis thaliana* and *Nicotiana benthamiana*, from *Phakopsora pachirhyzi* in *N. benthamiana* and from *Hyaloperonospora arabidopsidis* in *A. thaliana* allowed the study of their subcellular localization *in planta*, their impact on the growth of different pathogens and the search for host proteins potentially targeted by CEs [19, 25-27].

Still, the impact of CEs in the plant may not be easy to detect or the isolated effect of a single CE may be too subtle to affect pathogen growth. In the study of Germain and colleagues, 14 CEs impacted the growth of *H. arabidopsidis* or *Pseudomonas syringae* pv *tomato*. Eleven of the analyzed CEs displayed nucleocytoplasmic localization *in planta*, providing very limited information on possible host targets or helpers of these protein [19]. Petre and colleagues found seven CEs of wheat yellow rust fungus (out of 16) with specific accumulation pattern in plant cells (other than nucleocytoplasmic) and discovered specific plant protein interactors for six CEs [28]. Only three of the 16 CEs studied had both specific accumulation pattern in *N. benthamiana* cells and specific plant protein interactors. Although the pathogen growth readout is informative regarding the impairment of the immune pathway, it is opaque with regards to which pathway has been tampered with or which metabolites are off-balance. Transcriptomic and metabolomic studies of stable transgenic plants expressing CEs have been useful in these cases, since they allow the detection of more subtle changes, unlikely to have a quantifiable impact on pathogen growth on their own [22, 29, 30].

Here we studied the transcriptome and metabolome of 14 transgenic *Arabidopsis* plant lines expressing *Mlp* CEs known to affect plant susceptibility to pathogen. We identified 2 299 deregulated genes using this approach, including many related to response to biotic and abiotic stress, metabolism of specialized metabolites and plant development. Four lines expressing CEs from different families showed correlated patterns of gene deregulation demonstrating that the current grouping based on sequence homology does not reflect the virulence function of these CEs. We also found important down-regulation of highly unsaturated and phenolic compounds and up-regulation of peptides in almost all CE-overexpressing lines. Overall, our results show a lack of correlation between the sequence similarity of the studied CEs and their overall deregulation of genes or metabolites. Taken together, our results demonstrate that CEs that have completely different sequences can alter the expression of the same genes sets, while CEs of the same family can target completely different gene sets. Therefore, it is not possible to estimate the function of a CE, its impact on the transcriptome or on the metabolome of the plant, based solely on its sequence, or its similarity to another CE.

## Results

### *In planta* expression of candidate fungal effectors results in important deregulation at the transcriptome level

*Melampsora larici-populina* CEs have been previously studied in heterologous systems for functional characterization [19, 22, 25, 30, 31]. In Table 1, we present features of the 14 CEs studied here. Mlp37347 is a homolog of the well studied AvrL567 group from *M. lini* [32, 33], and accumulates at the plasmodesmata in *Arabidopsis*. Mlp72983 accumulates in the chloroplast [19] and Mlp124357 is found in the tonoplast and was shown to interact with *Arabidopsis* and poplar Protein Disulfide Isomerase [30]. The other 11 CEs selected here have nucleocytosolic accumulation, the same as the marker protein GFP used. Although information about these CEs is scarce, all of them impacted *Arabidopsis* susceptibility to either *Pseudomonas syringae* or to *Hyaloperonospora arabidopsidis*.

**Table 1.** Features of the CEs investigated in this study.

| CE | Length<br>(Cysteine) | Family (members) | Subcellular<br>localization<sup>a</sup> | U, P, B, L<sup>b,c</sup> |
| --- | --- | --- | --- | --- |
| Mlp37347 | 151 (2) | - | Plasmodesmata | E, HE, E, E |
| Mlp72983 | 220 (8) | CPG332-CPG333(13) | Chloroplast | E, HE, E, HE |
| Mlp102036 | 107 (0) | CPG2528(5) | Nucleocytosolic | E, HE, E, E |
| Mlp106078 | 137 (10) | - | Nucleocytosolic | E, HE, E, E |
| Mlp123218 | 209 (6) | CPG543(7) | Nucleocytosolic | E, HE, E, E |
| Mlp123227 | 124 (3) | CPG1059(2) | Nucleocytosolic | E, HE, E, HE |
| Mlp123531 | 102 (8) | CPG4557(3) | Nucleocytosolic | E, HE, E, E |
| Mlp124256 | 89 (6) | CPG5464(13) | Nucleocytosolic | N, N, E, E |
| Mlp124266 | 92 (7) | CPG5464(13) | Nucleocytosolic | N, N, E, E |
| Mlp124357 | 98 (6) | CPG4890 | Tonoplast | N, N, E, E |
| Mlp124466 | 76 (0) | - | Nucleocytosolic | - |
| Mlp124497 | 77 (4) | CPGH1(33) | Nucleocytosolic | N, N, N, N |
| Mlp124499 | 72 (3) | CPGH1(33) | Nucleocytosolic | N, N, E, HE |
| Mlp124518 | 76 (3) | CPGH1(33) | Nucleocytosolic | N, N, E, E |

<sup>a</sup> Subcellular localization was evaluated in Arabidopsis [19]. <sup>b, c</sup> U, P, B, L refers to expression on: U) urediniospores, P) poplar leaves, B) basidiospores or L) larch needles [46], where E, HE, and N indicate that the CE is expressed, highly expressed, or was not detected, respectively, and - indicates no data is available.

To better understand the mechanism through which these 14 CEs impact plants, we studied the transcriptome and the metabolome of transgenic *Arabidopsis* plants constitutively expressing them. In total, we found 2 299 differentially expressed genes (DEGs) across the experiment. However, the number of DEGs in each line was variable, from 84 in Mlp106078 to 898 DEGs in Mlp123531 (Fig 1), indicating each CE affects the plant transcriptome to a different degree. The list of deregulated genes in each transgenic line is available in Table S1. We further assessed if the level of transgene expression could explain the number of DEGs in each sample and plotted the number of deregulated genes per transgenic line against expression level (in transcript per million) of the CE:GFP fusion transcripts. Linear regression shows a poor relation between the two (R^2^ = 0.1016, Fig S1) suggesting that the number of deregulated genes per line depends more on the identity of the expressed CE than on the strength of its expression.

**Fig 1.**
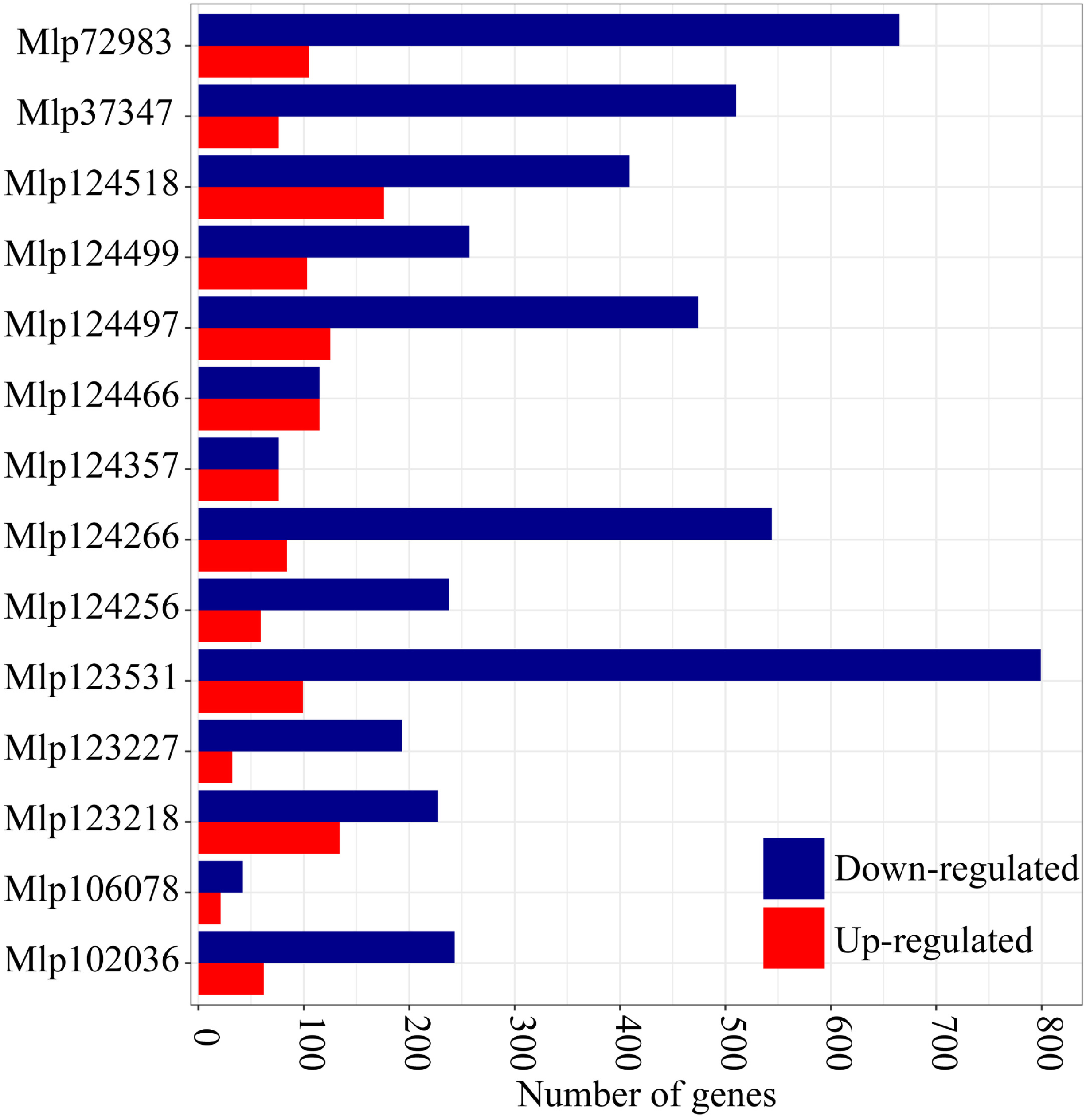
*In planta* expression of candidate fungal effector results in important deregulation at the transcriptome level. Blue and red bars indicate the number of down- and up-regulated genes, respectively, in each CE-expressing transgenic line compared to the control line. The underlying data for this figure can be found at dos Santos et al. [34].

### Hierarchical clustering based on gene expression groups effectors independently of amino acid sequence homology

CEs are typically grouped into families based on their amino acid sequences [11] and it has been shown that R-protein recognize related effectors [12]. Nevertheless, the virulence activity of effectors from the same family has rarely been studied. To search for gene deregulation patterns of related and unrelated CEs, we used WGCNA to cluster the co-expressed DEGs and Pearson’s correlation coefficient to cluster the transgenic lines (Fig 2). We found in total 208 GO terms enriched in the gene sets from WGCNA. A summary is presented in Table 2, and the full list of enriched terms is available at dos Santos et al. [34]. Set 0 clusters 714 genes deregulated across the 14 transgenic lines, 63.17% of which were down-regulated. Functions enriched in this gene set are related to defense, specialized metabolism, stress, and signaling pathways. Set 1 is composed of down-regulated genes enriched in GO terms related to defense responses and all transgenic lines have down-regulated genes in this set. Of the 379 genes in Set 2, 76.5% were down-regulated and this set is enriched GO terms related to specialized metabolite biosynthesis. In the case of Set 3, 81.8% of the genes were down-regulated, but we did not find enriched GO terms in this gene set. Interestingly, this set is composed of genes with the same pattern of deregulation in 4 transgenic lines expressing effectors without sequence similarity (Mlp72983, Mlp102036, Mlp123218, and Mlp123531, Table S2) which accumulate in two separate cell compartments (Table 1). Set 4 is related to metabolism and abiotic stress and 77.6% of its genes were down-regulated. Sets 5, 6, and 7 are composed almost exclusively of up-regulated genes (Table 2). Set 5 has genes deregulated in most transgenic lines that are related to abiotic stress and development. Set 6 is comprised of up-regulated genes almost exclusively found in the transgenic line Mlp124466 and related to transcription, vascular histogenesis, and response to different types of stress. Finally, Set 7 is made of genes related to photosynthesis and deregulated in the lines Mlp124256 and Mlp124518. In the cases of the Sets 0, 2, 3 and 4, there is mix of genes up and down-regulated, thus the enriched GO terms may be either up or down-regulated, or both. Interestingly, the dendrogram at the top of Fig 2 shows that CEs belonging to the same family (Mlp124497, Mlp124499 and Mlp124518; Mlp124256 and Mlp124266) fall in separate clusters despite their similarity at the amino acid level (Table S2).

**Table 2.** Summary of “biological process” GO terms enriched in the WGCNA gene sets.

| Set | Genes in the<br>set | Up-regulated<sup>a</sup> | Down-regulated<sup>a</sup> | Enriched GO terms |
| --- | --- | --- | --- | --- |
| Set 0 | 714 | 262 | 451 | Response to water deprivation<br>Cold acclimation; Leaf senescence<br>Response to fungus, to chitin, to ROS<br>Response to salt stress and to hypoxia<br>Defense response to fungus<br>Response to toxic substance<br>Response to nitrogen compound and to ET |
|  |  |  |  | Isoprenoid, triterpenoid and terpenoid biosynthesis |
|  |  |  |  | Plant-type cell wall loosening |
|  |  |  |  | Phosphorelay signal transduction system |
| Set 1 | 624 | 10 | 615 | Response to drug, nitrogen, ROS and ozone |
|  |  |  |  | Response to SA, JA and karrikin |
|  |  |  |  | Response to wounding, to herbivore and insect |
|  |  |  |  | Cellular response to light stimulus and hypoxia |
|  |  |  |  | Cellular response to acid chemical |
|  |  |  |  | Defense response (incompatible interaction) |
|  |  |  |  | Defense response by callose deposition in cell wall |
|  |  |  |  | Defense response by cell wall thickening |
|  |  |  |  | SAR and ISR |
|  |  |  |  | Camalexin, indole phytoalexin and SA biosynthesis |
|  |  |  |  | Sulfur compound biosynthesis |
|  |  |  |  | Toxin and phenol-containing compound biosynthesis |
| Set 2 | 379 | 89 | 290 | Response to karrikin, to nutrient levels and to copper ion |
|  |  |  |  | S-glycoside and unsaturated fatty acid biosynthesis |
|  |  |  |  | Chlorophyll biosynthesis |
|  |  |  |  | Tetraterpenoid, terpenoid and carotenoid biosynthesis |
|  |  |  |  | Isoprenoid, glycosyl and xanthophyll metabolism |
|  |  |  |  | Sulfur compound, cofactor and leucine biosynthesis |
|  |  |  |  | Defense response to insect |
|  |  |  |  | De-etiolation; Chloroplast organization |
| Set 3 | 253 | 47 | 207 | No GO term enriched |
| Set 4 | 140 | 32 | 109 | Response to water deprivation |
|  |  |  |  | Response to salt stress and to starvation |
|  |  |  |  | Cellular amino acid catabolism/metabolism |
|  |  |  |  | ET-activated signaling pathway |
|  |  |  |  | Indole-containing compound metabolism |
| Set 5 | 116 | 113 | 4 | Circadian rhythm; Starch catabolism |
|  |  |  |  | Response to cold |
|  |  |  |  | Regulation of reproductive process |
|  |  |  |  | Regulation of post-embryonic development |
| Set 6 | 40 | 38 | 2 | Response to hypoxia and to wounding, |
|  |  |  |  | Response to drug, to chitin and to salt stress |
|  |  |  |  | Transcription; Phloem or xylem histogenesis |
| Set 7 | 32 | 32 | 0 | Photosynthesis; Proton transmembrane transport |

<sup>a</sup> Up- and down-regulated indicate the number of genes in the set the are up- or down-regulated in at least one transgenic line, thus there may be genes that are deregulated in both directions in the set because they are deregulated in opposite directions in different samples.

**Fig 2.**
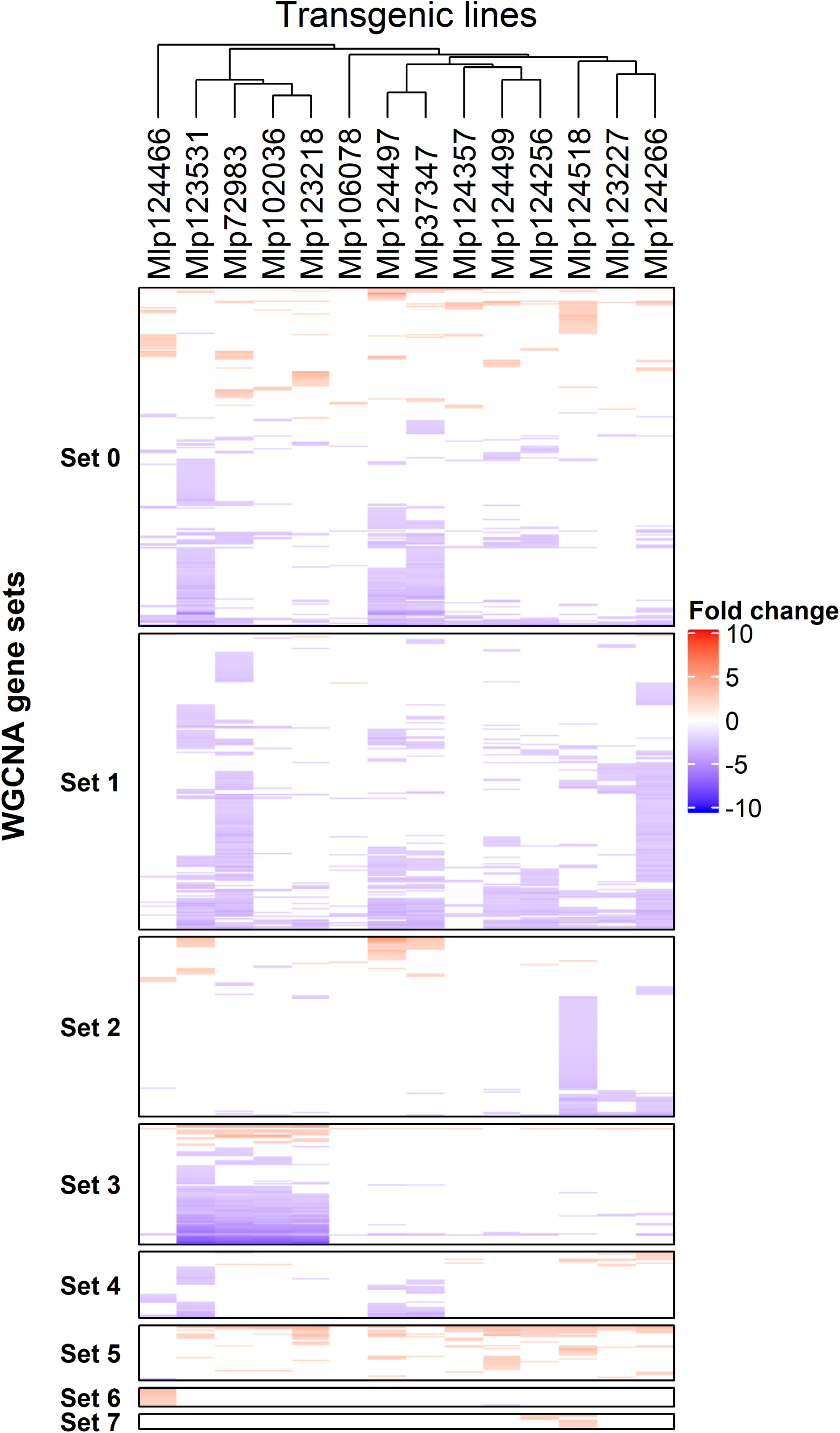
Heatmap of genes deregulated in each CE-expressing transgenic line. Transgenic lines are displayed as columns and deregulated genes as lines. Sets of co-expressed genes (Sets 0 to 7) were calculated with WGCNA. Transgenic lines were grouped by correlation of gene deregulation using Pearson’s correlation coefficient. The underlying data for this figure can be found at dos Santos et al. [34].

To analyze the relation between the sequence of each effector and its influence on the plant transcriptome, we compared the sequence alignment dendrogram to the differential expression dendrogram. After removal of the signal peptide, we aligned the sequences of the studied CEs, and compared the resulting dendrogram with the one obtained from the gene deregulation correlation (Fig 3). Pearson’s correlation showed that transgenic lines expressing CEs from different families had correlated patterns of gene deregulation. Only one cluster was present in both dendrograms, Mlp102036 and Mlp123218, however this grouping is not supported in the effector sequence dendrogram (bootstrap value 8%) while it is in the gene deregulation dendrogram (bootstrap 100%). This analysis indicates that the sequence similarity between the CEs is not a good predictor of the impact they have on plant gene expression.

**Fig 3.**
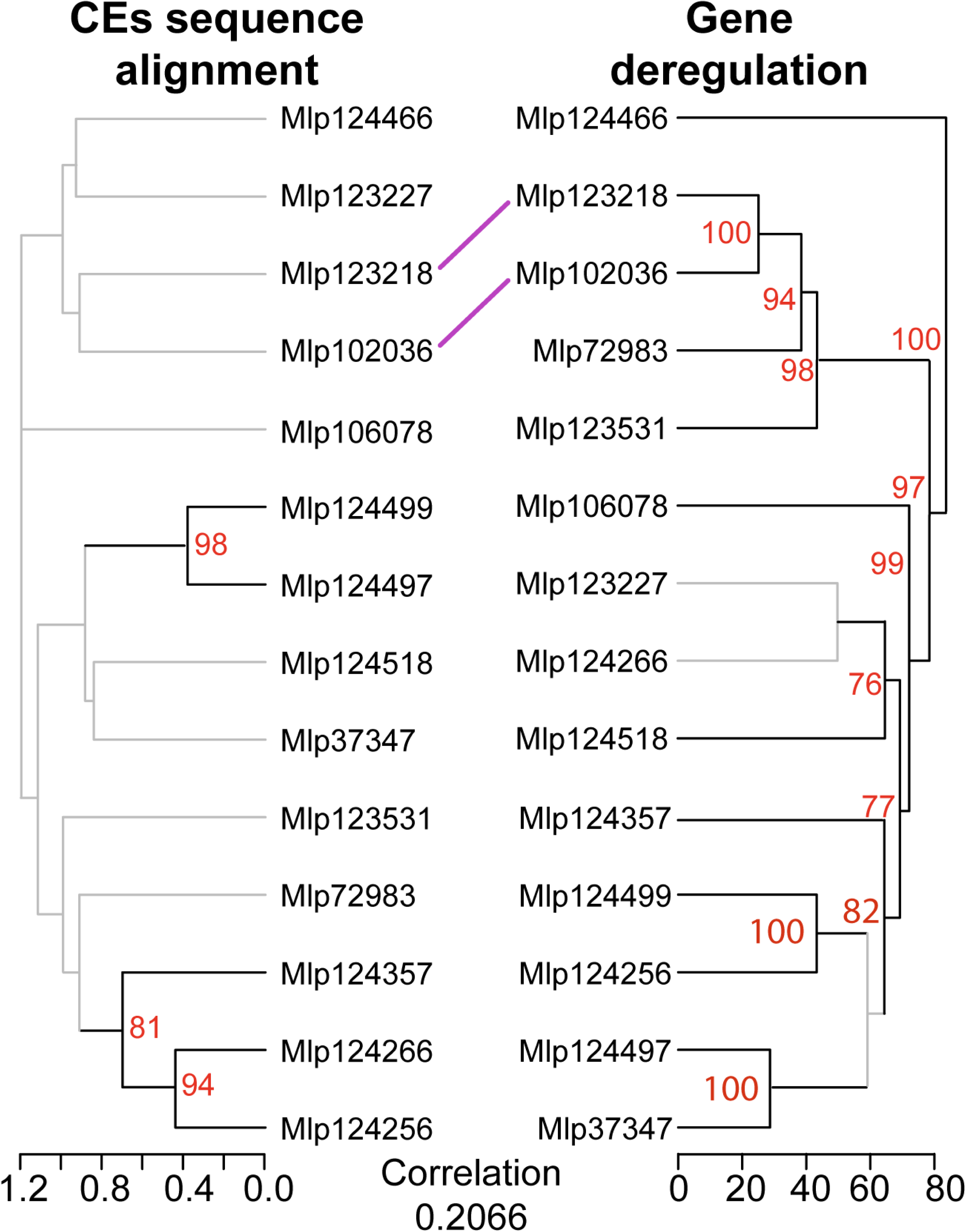
Hierarchical clustering of gene deregulation groups effectors independently of amino acid sequence homology. CE sequence alignment was computed with Muscle alignment and tree (left) was calculated with UPGMA. Dendrogram based on correlation of gene deregulation (right) was calculated with Pearson’s correlation coefficient of Fold Change levels and bootstrap values were obtained with pvclust. Branches with bootstrap support < 70% are shown in grey. Central lines indicate shared clusters and cophenetic correlation between the dendrograms is shown in the bottom. The underlying data for this figure can be found at dos Santos et al. [34].

### Effectors converge on deregulating the same metabolic pathways while others display unique patterns

Even though the transcripts affected by related effectors are different, in theory they could fall within the same metabolic pathway and therefore similarly alter the plant. To test this hypothesis, we searched for KEGG pathways over-represented in the up- and down-regulated genes in each transgenic line. “Biosynthesis of secondary metabolites” and “Metabolic pathways” were enriched among gene sets (either up-, red, or down-regulated, blue) of eight transgenic lines, while “MAPK signaling pathway” and “Plant-pathogen interaction” were enriched only among the down-regulated genes of six and five transgenic lines, respectively (Fig 4). We also found that “Starch and sucrose metabolism” was down-regulated in the transgenic lines Mlp123227 and Mlp124266, but up-regulated in the lines Mlp123218 and Mlp124497, whereas several transgenic lines showed impact on specialized metabolism. This was also visible in the enriched GO terms found on the WGCNA gene sets (Table 2 and [34]). File S1 shows heatmaps of 11 different metabolic pathways in which there were at least 10 genes deregulated across the experiment. The circadian rhythm pathway, although enriched only among the down-regulated genes of the lines Mlp124499, Mlp37347 and Mlp123531 and up-regulated genes in the Mlp124357 transgenic line, has several genes deregulated in all the transgenic lines studied. The plant-hormone signal transduction pathway is enriched among down-regulated genes in the transgenic lines Mlp37347, Mlp123531, and Mlp124497, and which we found several down-regulated genes (17, 23, and 17 DEGs, respectively) related to auxin response. From these results we conclude that CEs with similar sequences not only deregulate different genes but also alter different pathways.

**Fig 4.**
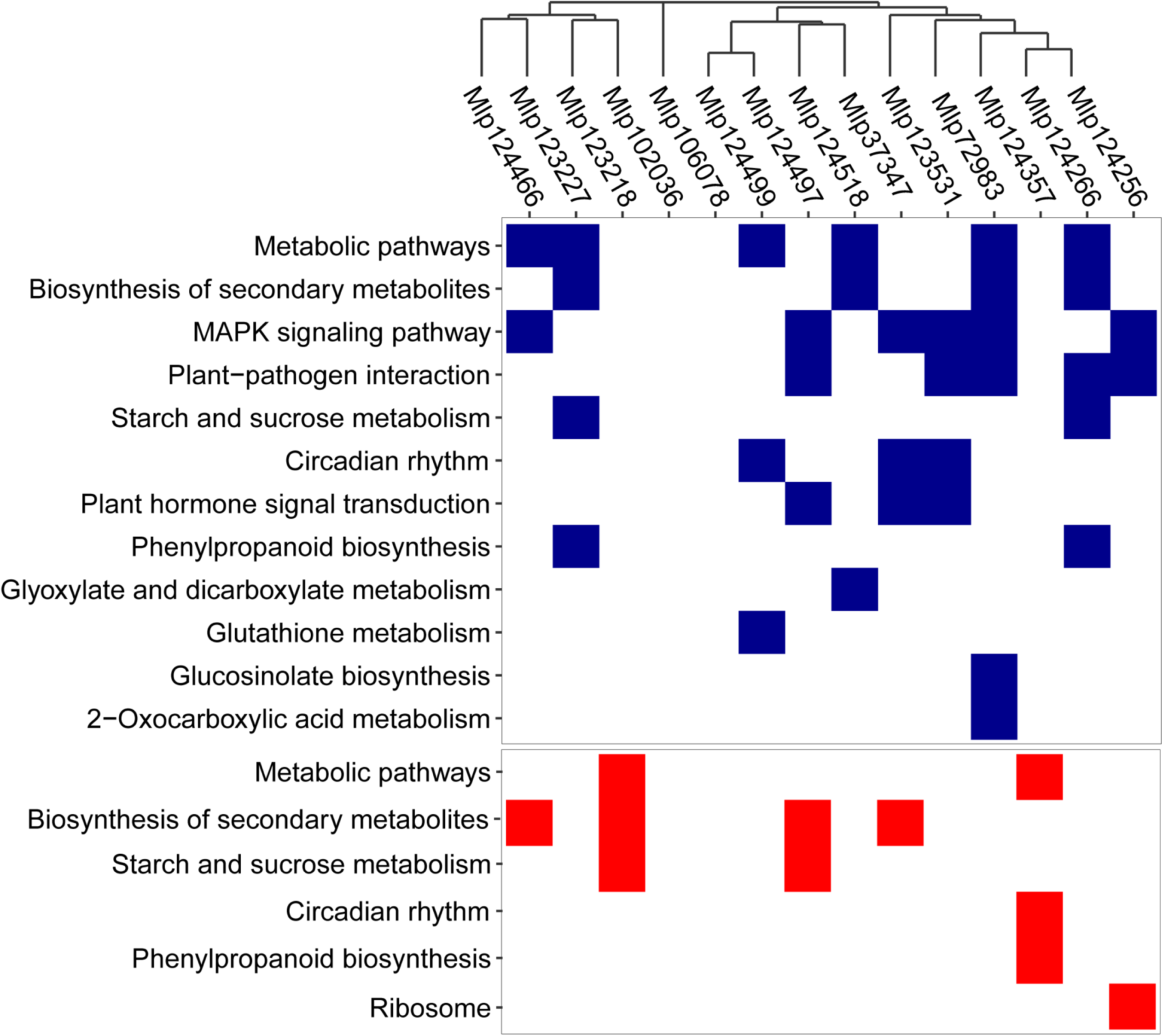
Effectors converge on deregulating the same metabolic pathways while others display unique patterns. KEGG pathways over-represented among the sets of genes down-(blue) and up-regulated (red) in each transgenic line (columns) were calculated with KEGGprofile. Transgenic lines are ordered according to dendrogram of sequence similarity calculated with Muscle. The underlying data for this figure can be found at dos Santos et al. [34].

As both primary and specialized metabolisms were affected at the transcriptomic level and their levels can have an important role in the outcome of an infection, we proceeded with an untargeted analysis of the metabolome of these plants. We extracted metabolites with solutions containing 20% and 80% methanol and used ultra-high resolution mass spectrometry in negative mode. A total of 5 192 masses were assigned across the experiment, ranging from 2 679 (Mlp123227) to 3 151 (Mlp124357) masses in each transgenic line (Table S3). When separated in biochemical categories, assigned formula belonged mostly to highly unsaturated and phenolic and aliphatic categories, while peptides, sugars, condensed aromatics and polyphenolics were less important both in number of formulas and in relative abundance (Fig 5). Compared to the control, we found 680 assigned molecular formulas with a | log_2_-transformed Fold change | > 2 (Fig 6A), ranging from 69 metabolites in the line Mlp124466 (1.95% of the masses detected in this sample and/or in the control) to 353 in the line Mlp123227 (9.68% of the masses detected in this line and/or in the control, Table S3). In all transgenic lines, with exception of Mlp72983 and Mlp124256, there was over-representation of highly unsaturated and phenolic compounds among the down-regulated metabolites (accumulation level lower than in the control line) whereas up-regulated metabolites (accumulation level higher than in the control line) were enriched in peptides in all samples, except Mlp72983, Mlp106078 and Mlp124466 (Fig 6B, Table S4). As done with the transcriptomic data, we assessed whether the variation in the number of metabolites deregulated in each transgenic line could be explained by the level of expression of the transgene. For this, we plotted the number of deregulated metabolites per transgenic line (left Y-axis, blue) against the average expression level of the CEs in each transgenic line (X-axis, Fig S2). As the number of metabolites detected in each transgenic line varied (Fig 5A), we also plotted the ratio of deregulated metabolites:identified (detected either in the control or in the corresponding sample) metabolites in the right Y-axis (red). We found that the variation in transgene expression could explain neither the number (R^2^=0.0063, p-Value=0.7872) nor the ratio of deregulated metabolites (R^2^=0.0033, p-Value=0.8444), suggesting that the magnitude of the impact on the metabolome depends on the identity of the CE expressed in the plant rather than the strength of the CE expression.

**Fig 5.**
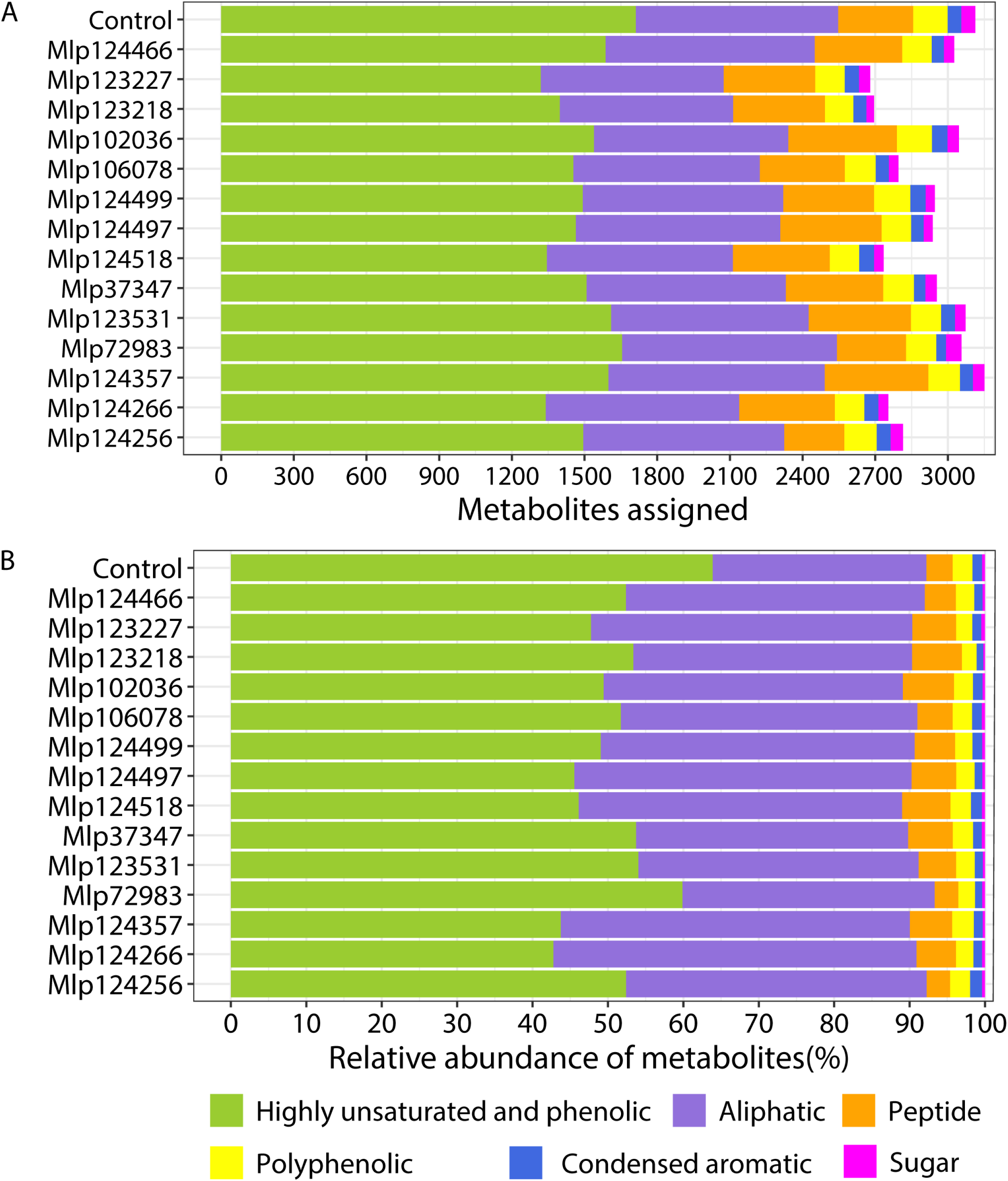
Metabolic composition of samples in number of formulas (A) and relative abundance of compounds (B). Samples were analyzed in negative mode and estimated molecular formulas were separated in six categories: highly unsaturated and phenolic (green), aliphatic (purple), peptide (orange), polyphenolic (yellow), condensed aromatic (blue), and sugar (pink). The underlying data for this figure can be found at dos Santos et al. [34].

**Fig 6.**
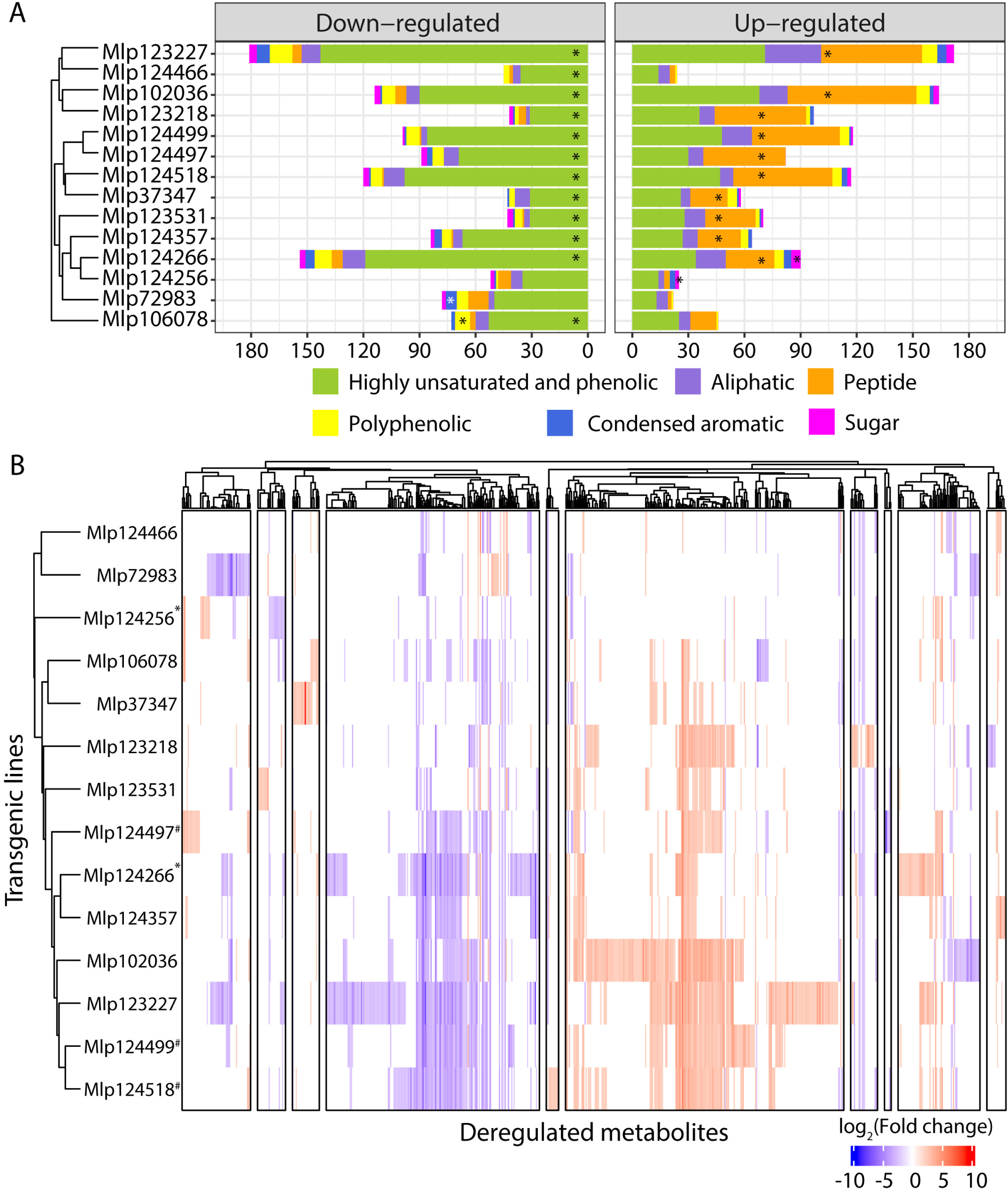
(A) Metabolites down-regulated (left) are enriched in highly unsaturated and phenolic compounds while peptides are over-represented among those up-regulated (right). Samples were analyzed in negative mode and relative abundance of metabolites in samples was compared to that in the control plants. Estimated molecular formulas were separated in six categories: highly unsaturated and phenolic (green), aliphatic (purple), peptide (orange), polyphenolic (yellow), condensed aromatic (blue), and sugar (pink). (B) Transgenic lines expressing candidate effectors with no similarity in amino acid sequence have correlated patterns of metabolite deregulation. Both metabolites and transgenic lines were clustered using Pearson’s correlation. ^*^ indicates transgenic lines with CEs from the CPG5464 family; ^#^ indicates transgenic lines with CEs from the CPGH1 family. The underlying data for this figure can be found at dos Santos et al. [34].

In order to find shared patterns of metabolite deregulation across the transgenic lines studied, we used Pearson’s correlation to group metabolites with correlated deregulation across the experiment and transgenic lines which deregulated the same metabolites. As observed with the gene deregulation, we found that transgenic lines expressing CEs without sequence similarity have correlated patterns of metabolite deregulation (Fig 6B). In the case of the CPGH1 family (CEs Mlp12497, Mlp124499, Mlp124518), lines Mlp124499 and Mlp124518 are correlated at 0.77 (Pearson’s correlation), but their correlation with the line Mlp124497 is less strong (Mlp12497-Mlp124499: 0.59; Mlp124497-Mlp124518: 0.64). The two AvrP4 homologues, Mlp124256 and Mlp124266, have 46.3% of amino acid sequence similarity [34], but the correlation in metabolites deregulation patterns of the transgenic lines expressing these CEs is of 0.32. On the other hand, although Mlp124266 and Mlp124357 have 21.2% of amino acid sequence similarity (Table S2), multiple sequence alignment groups the AvrP4 homologues with the CE Mlp124357 (Fig 7) and their metabolite deregulation correlation is 0.69.

**Fig 7.**
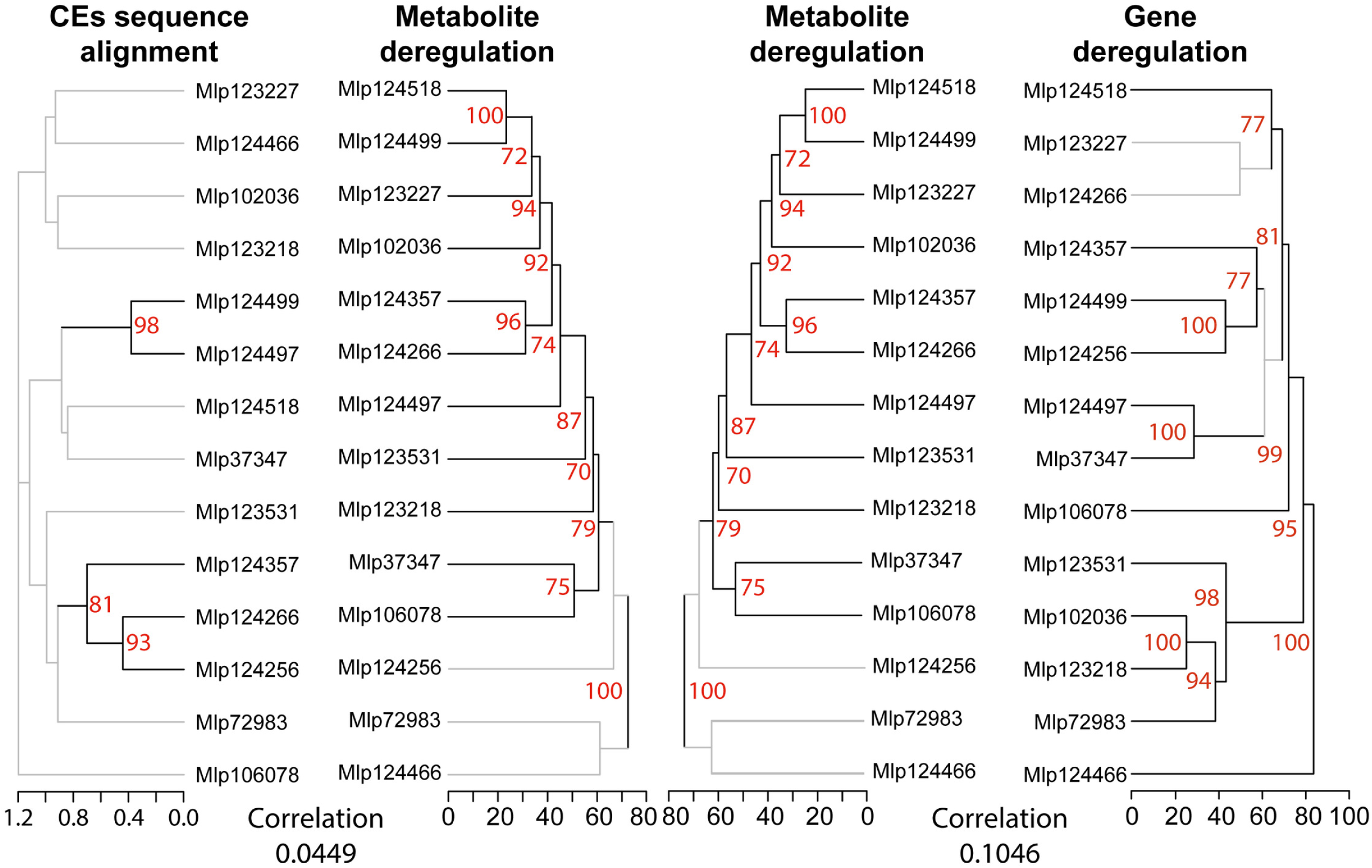
Hierarchical clustering based on metabolite deregulation groups effectors independently of amino acid sequence homology and gene deregulation patterns are not correlated to metabolite deregulation patterns in CE-expressing lines. CE sequence alignment was computed with Muscle alignment and tree (left) was calculated with UPGMA. Dendrograms based on correlation of metabolite deregulation (center) or gene deregulation (right) were calculated with Pearson’s correlation coefficient of Fold Change levels and bootstrap values were obtained with pvclust. Branches with bootstrap support < 70% are shown in grey. The underlying data for this figure can be found at dos Santos et al. [34].

Remarkably, there was no correlation between the dendrograms gene and metabolite deregulation (cophenetic correlation of 0.1046, Fig 7). When considering the number of genes and metabolites deregulated in each sample, the correlation was also low (Pearson’s correlation = −0.1182). These results suggest these two omics approaches are needed to understand the magnitude of the impact of the CEs in the plant. Nevertheless, the possibility that the metabolic pathways deregulated at the metabolite level are the same as those deregulated in the gene level cannot be discarded.

To identify the molecular formula assigned in each sample and associate the metabolomic results with metabolic pathways, we searched for compounds with matching formula or matching *m/z* values in the KEGG database. From the 5 192 *m/z* detected across the experiment, 385 (7.42%) had a single match in KEGG database, while other 600 corresponded to multiple metabolites. When only considering the 680 deregulated metabolites, 54 (7.07%) matched a single metabolite and 82 (12.06%) matched multiple metabolites [34], leaving 546 unmatched.

## Discussion

Effector biologists have tackled both the identification and the functional characterization of candidate effectors (CEs) [14, 35], as this is a key step towards a better understanding of plant-microbe interactions. In rust fungi, different approaches are used in the functional characterization of these proteins, including analysis of subcellular localization *in planta* [19, 25, 28, 30, 31], infection assays in true host or in a model plant, and induction/repression of plant cell death [19, 30, 36, 37]. The transcriptome or metabolome of the host in responses to the pathogen are frequently evaluated [38-43], but the assessment of the role of individual CEs in these processes is not easily measured and seldom analyzed [22, 44]. Here we investigated 14 CEs from *Melampsora larici-populina* by evaluating their individual impact on the transcriptome and metabolome of stable transgenic *Arabidopsis* plants. By studying the impact of several individual CEs, we were able to compare patterns of gene and metabolite deregulation. Unexpectedly, we found that transgenic lines expressing Ces belonging to the same family did not have comparable patterns of gene or metabolite deregulation.

Previous studies in *M. larici-populina* have shown that genes encoding fungal effectors are expressed in waves in the telial host [45] and that members of the same family may be expressed during the infection of different hosts [46]. This reflects the functional diversification of effectors, indicating that the fungus uses different sets of effectors for each stage of the infection and suggesting that effector families can have different functions, may target different host proteins or the same host protein that diverged in different hosts. The concurrent study of individual *M. larici-populina* CEs allows the comparison of their individual impact in the plant [19]. We found variability in the magnitude of the impact of each CE on the transcriptome (from 84 to 898 DEGs) and the metabolome (from 69 to 363 metabolites deregulated, Figs 1 and 6) of the transgenic plants, a variability which is not related to the level of expression of the transgenes (Figs S1 and S2). This suggests that the identities of the CEs are orienting the deregulations. By comparing the correlation of gene and metabolite deregulation patterns with the CEs sequence similarity (Figs 3 and 7), we show that CEs belonging to the same family do not deregulate the transcriptome or the metabolome in a same way nor do they deregulate the same metabolic pathways (Fig 4). These results corroborate the infection assays from Germain and colleagues [19]. In their study, *Arabidopsis* plants, constitutively expressing *Mlp* CEs, were infected with *P. syringae* DC3000 or *H. arabidopsidis* Noco2. Mlp124497, Mlp124499 and Mlp124518 (family CPGH1) and Mlp124256 and Mlp124266 (family CPG5464) [47], all increased *Arabidopsis* susceptibility to *H. arabidopsidis*. However, only Mlp124266, Mlp124497 and Mlp124499 made *Arabidopsis* more susceptible to *P. syringae*.

It has been suggested that proteins with higher sequence similarity have higher probability of having the same function [48], thus small secreted proteins from many fungal and oomycete plant pathogens [9, 11, 49-51] have been grouped in protein families to guide functional annotation and to help understand effector evolution. Nevertheless, recent studies have hypothesized that effectors from the same family may have different functions in the same host. This is the case for HopAF1 effectors from *P. savastanoi* [52] and GALA effectors from *Ralstonia solanacearum* [53], which impact differently the plant defense. It is also the case for XopD effectors from plant pathogenic bacteria, which show different levels of SUMO protease activity and have different impacts in *Nicotiana* leaves [54]. This hypothesis is also supported by the evolution of the Tin2 effector in Ustilaginaceae. Tin2 from *Ustilago maydis* interacts with *Zea mays* TTK1 protein to stabilize it, leading to accumulation of anthocyanin. However, Tin2 from *Sporosorium reiliannum* interacts with *Zea mays* TTK2 and TTK3, inhibiting their activity [55].

The CEs studied here deregulate diverse biochemical pathways in the plant (Fig 4). In relation to primary metabolism, genes in the “starch and sucrose metabolism” pathway were over-represented among up-regulated genes in the transgenic lines expressing the CEs Mlp123218 and Mlp124497, comparable to what is observed in susceptible wheat infected with *Puccinia triticina* [43]. On the other hand, the plants expressing Mlp123227 and Mlp124266 showed an enrichment of this pathway among down-regulated genes and the transgenic lines Mlp72983 and Mlp124266 had several genes down-regulated in this pathway as well (File S1), a pattern seen in resistant wheat infected with *P. triticina* [43]. This difference in the direction of gene deregulation within the same pathway by different CEs may be an indication that deregulated genes have different functions. It can also suggest that these CEs are used in different stages of the infection. When considering pathways related to defense, the transcriptomic deregulations found in this study differ from previous reports of susceptible plants infected by rust fungi. While genes encoding Glutathione-S-transferase are down-regulated in at least one of 12 transgenic lines studied here (File S1), these genes are up-regulated in apple leaves infected with *Gymnosporangium yamadae* [39]. Moreover, Tremblay and colleagues [40] reported up-regulation of genes in the “photosystem” and “nitrogen metabolism” pathways in susceptible *Glycine max* infected with *P. pachyrhizi*, whereas genes from these pathways were down-regulated in our transgenic lines.

There are several possible explanations for the differences between previous studies and our own. First, our results may be due to the long-term exposure of our plants to CEs, as they are stable transgenic lines, whereas during the infection rust fungi secrete effectors in waves [45], these proteins are not constitutively present in the host. It is also possible that results from Tao and colleagues [39] and Tremblay and colleagues [40] included the activation of PTI as well as the combinatory effect of multiple effectors, as they investigated plant response to the fungal infection, not to individual CEs. Our approach was to express CEs from *M. larici-populina* in a plant that cannot be infected by this fungus, thus should not recognize these proteins nor mount active defense responses against them. Nevertheless, as we evaluated the impact of each CE in the plant using one single transgenic line, it is not possible to know for sure if the impact on the transcriptome and metabolome is caused by the CE or is a secondary effect of the DNA insertion site in each of these transgenic lines. Yet, the probability that the insertion site impacted in the same manner the results of all the 14 transgenic lines studied here is low, thus the results that consider the 14 transgenic lines are robust. Finally, although there are limitations in the use of heterologous systems, they allow faster functional characterization of CEs [17, 56] and they may be indispensable for high-throughput studies of CEs of obligate biotrophic pathogens or other microorganisms not amenable to genetic manipulation [57, 58].

Taken together, our results reinforces the hypothesis that the CEs studied here and functionally characterized by Germain and colleagues [19] are *bona fide* effectors. Nevertheless, future studies interested in CEs evaluated here should analyze more independent transgenic lines. In addition, since our methodology for the metabolomic analysis is semi-quantitative and does not allow the distinction of metabolites with the same *m/z*, follow up studies should use chromatography in tandem with mass spectrometry and should analyze more replicates for the mass spectrometry. Our study also questions the validity of grouping CEs by sequence similarity. The importance of this approach for understanding the evolution of effectors is obvious [9], but basing functional characterization on sequence similarity may be misleading [52, 53, 55].

## Materials and Methods

### Plant growth conditions

*Arabidopsis thaliana* transgenic plants in Columbia-0 background expressing GFP alone (control) or fused to a candidate effector of the fungus *Melampsora larici-populina* (Mlp37347, Mlp72983, Mlp102036, Mlp106078, Mlp123218, Mlp123227, Mlp123531, Mlp124256, Mlp124266, Mlp124357, Mlp124466, Mlp124497, Mlp124499, Mlp124518) previously obtained in our laboratory [19, 30], were grown at 22°C at 12h/12h light/dark cycles.

### RNA extraction and transcriptome analysis

RNA was extracted from pooled aerial tissue of 2-week-old soil-grown plants, using three replicates per genotype, with the Plant Total RNA Mini Kit (Geneaid) using RB buffer following manufacturer’s protocol. The samples were treated with DNAse, then RNA quality was assessed using agarose gel electrophoresis. QC was performed using a 2100 Bioanalyzer (Agilent) and only samples having an RNA Integrity Number higher than 7 were kept for library preparation. Libraries were generated with the NeoPrep Library Prep System (Illumina) using the TruSeq Stranded mRNA Library Prep kit (Illumina) and 100 ng of total RNA as per the manufacturer’s recommendations. The libraries were then sequenced with Illumina HiSeq 4000 Sequencer with paired-end reads of 100 nt at the Genome Quebec Innovation Centre (McGill University, Montreal, Canada).

The bioinformatic analyses were done with Compute Canada servers, the parameters used are presented in Table S5. We trimmed the reads using Trimmomatic [59] and we aligned the surviving paired reads to the genome of *A. thaliana* assembly TAIR10 with HISAT2 [60]. Unmapped reads were aligned to the sequences of the CEs, without signal peptide, attached to eGFP. We counted the reads assigned to each transcript with the R (v3.6) packages Rsamtools (v2.2.3 [61]), GenomicAlignments and GenomicFeatures [62]. The general information of the sequencing results and mapping data is presented in Table S6. Before comparing the samples, we used the CustomSelection package [63] to select as reference genes the top 5% genes with lowest coefficient of variation of TPM among the 45 samples [34]. We assessed the variation between the replicates and the similarity of the samples with principal component analysis (Fig S3). Differential expression analysis was performed with DeSeq2 [64], using the un-normalized counts as input, and genes with |log_2_ Fold change| ≥ 2 (p-Value ≤ 0.01), when comparing each CE-expressing lines to the control line, were considered as deregulated. We used clusterProfiler[65] for GO term enrichment analysis and KEGGprofile (v1.24.0 [66]) for KEGG enrichment analysis. Sets of deregulated genes were computed using WGCNA [67]. We calculated the similarity of gene deregulation of different transgenic lines with the R package pvclust (v2.2-0 [68]), using Pearson’s correlation and 5 000 bootstrap replications.

### Metabolite extraction and metabolomics analysis

Metabolites were extracted from pooled aerial tissue of 2-week-old soil-grown plants, with four replicates per genotype. After pulverizing the tissues with a TissueLyser (30 cycles per second for 45 seconds repeated 3 times), we added 300 μL of distilled water to it. From the mix of tissue and water, we used 100 μL of tissue slurry for an extraction with 1 mL of 20% methanol and a separate 100 μL for an extraction with 1 mL of 80% methanol. After agitation with the solvent, we pooled the samples of the same genotype and extraction together and filtered them using glass microfiber filters (Whatman GF/F CAT No. 1825-025). We evaporated the extracts with a speed vacuum at room temperature and chamber vacuum of 7.4 torr’s and resolubilized them in 2 mL of distilled water. Then, we solid phase extracted 50 µg of dissolved organic carbon (DOC) of each sample, using Agilent PPL cartridges, and eluted it in 1mL of 100% methanol.

The mass spectrometry was performed in an Orbitrap LTQ-Velos calibrated and tuned to maximize the peak at 369.1 in Suwannee River Fulvic Acid (SRFA) reference material. The extracts were analysed by direct injection in negative mode at a resolution setting 100 000, with accumulation time set to a maximum of 500 ms and a target of 1 x 10^6^ ions. Peaks were only considered for formula assignment if their intensity was higher than 10x the median noise baseline. We assigned formulas to masses using an in-house MATLAB script [69] and we allowed assignments with mass error < 2 ppm. Briefly, formulas were considered over the ranges CH_4-100_O_2-40_N_0-2_ under the conditions O ≤ C; 0.3C ≤ H ≤ 2.2C. For each sample, the intensity of the peaks was normalised so that the sum of the intensities equalled 10 000.

Following analyses were performed using R software (v4.0). We used the molecular formulas to calculate the modified aromaticity index (AImod) of each metabolite [70] and the compound categories were defined as: condensed aromatic (AImod > 0.66), polyphenolic (0.66 ≥ AImod > 0.5), highly unsaturated and phenolic (AImod < 0.5 and H/C < 1.5), aliphatic (2 ≥ H/C ≥ 1.5, N= 0), peptide (2 ≥ H/C ≥ 1.5, N > 0) or sugar (O/C > 0.9) [71].

The results of the two extractions, with 20% and 80% methanol, were combined and the fold changes (FC) were calculated as 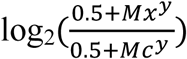 where 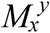 is the relative abundance of the metabolite *y* in the CE-sample *x* and 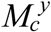 is the relative abundance of the metabolite y in the control. For each sample, only metabolites with |FC| > 2 were considered to have relative abundance different to that of the control. Categories enriched among up- and down-regulated genes were found by applying Fisher’s test. We calculated the similarity of metabolite deregulation of different transgenic lines with the R package pvclust (v2.2-0 [68]), using Pearson’s correlation and 5 000 bootstrap replications. Pairwise correlation of metabolite deregulation between specific transgenic lines was calculated with the function *cor* from the R package stats, using the method “pearson”. We were not able to analyze the extraction with 80% methanol of the transgenic line Mlp123218, thus the results presented for this line are only of the extraction with 20% methanol and they are compared to the results of the Control for the same extraction for consistency.

We searched the molecular formulas, obtained with the in-house script, in KEGG database using the R package KEGGREST (version 1.24.0 [66]) for identification of the metabolites detected. We also used Pathos [72] to search for metabolites with the same *m/z* (settings: negative mode, all organisms, -H^+^ as adduct and mass error at 3 ppm).

### Sequence analysis and integration

Multiple sequence alignment of CE amino acid sequences without signal peptides was performed with the software MEGA X [73] using Muscle [74] default settings. Evolutionary history was inferred using UPGMA method and 1 000 bootstrap replicates. Comparisons of dendrograms from CE sequence alignment, gene and metabolite deregulation correlation were done with dendextend R package [75] by calculating the cophenetic correlation between two dendrograms. We performed pairwise sequence alignment of the 14 CEs using Needle [76], with default parameters.

### Data availability

#### Transcriptomics

Raw reads and count matrices are available in NCBI GEO under the accession GSE158410 [77].

#### Metabolomics

Raw and mzXML files along with annotation of metabolites and their relative abundances in each sample are available at MetaboLights under the accession MTBLS2096 [78].

Data underlying figures full list of enriched GO terms in the WGNCA gene sets, information on the deregulated metabolites and the list of selected reference genes is available at [34].

## Acknowledgements

We thank Melodie B. Plourde and Benjamin Petre for critical review of the manuscript. Funding for the project was provided by Natural Sciences and Engineering Research Council of Canada (NSERC) Discovery Grants to HG. The project in HG’s laboratory was also partially funded by an institutional Research Chair and a Canada Research Chair held by HG and a Canada Research Chair held by IDP. KCGS was funded by a master’s scholarship from the Fondation de l’Université du Québec à Trois-Rivières, an international PhD scholarship from the Fonds de Recherche du Québec sur la Nature et les Technologies (FRQNT) and a graduate fellowship from MITACS.

---

**Fig S1. Magnitude of impact of CE on the plant’s transcriptome is independent of its level of expression**. Reads not mapped to *Arabidopsis* genome were aligned to the transgene sequences (CE:GFP fusion) and average expression (in transcripts per million) across replicates of each transgenic line was calculated. Linear regression was performed using the number of genes deregulated in each transgenic line as the dependent variable and the average expression of the CE as the independent variable. The underlying data for this figure can be found at dos Santos et al. [34].

**Fig S2. Magnitude of impact of CE on the plant’s metabolome is independent of its level of expression, considering either the absolute number of deregulated metabolites (triangles, linear regression results in blue) or the ratio of metabolites deregulated by those identified (circles, linear regression results in red)**. Reads not mapped to *Arabidopsis* genome were aligned to the transgene sequences (CE:GFP fusion) and average expression (in transcripts per million) across replicates of each transgenic line was calculated. Two separate linear regressions were performed using the number of metabolites deregulated and the ratio between metabolites deregulated by those detected in each transgenic line as the dependent variables and the average expression of the CE as the independent variable in both cases. The underlying data for this figure can be found at dos Santos et al. [34].

**Fig S3. Principal component analysis of the replicates of 14 transgenic lines expressing candidate effectors from *Melampsora larici-populina* attached to GFP and a control line expressing only GFP (black dots)**. Replicates of the same transgenic lines are close together, indicating the homogeneity of the sample, with exception of one replicate of each of the following transgenic lines: Mlp102036 (yellow), Mlp106078 (red), Mlp124256 (sky blue) and Mlp124357 (dark green). The underlying data for this figure can be found at dos Santos et al. [34].

**File S1. Heatmaps of deregulated genes separated by KEGG pathway**. Pathways with at least 10 genes deregulated across the experiment were selected for display of genes deregulated in each transgenic line. The underlying data for this figure can be found at dos Santos et al. [34].

**Table S1. List of deregulated genes across the experiment with log**_**2**_**-transformed fold changes (FC) and false discovery rates (FDR) for each transgenic line**.

**Table S2. Percentage of identity and similarity, presented as “ID (SIM)”, calculated with pairwise sequence alignment of CEs using Needle**.

**Table S3. Summary of metabolomic analysis in negative mode of extractions with 20% and 80% methanol combined**. Assigned, CHO, CHON and Mean mass refer exclusively to the sample in question, while the amount of deregulated formulas considers those *m/*z detected in the sample or in the Control.

**Table S4. Metabolites assigned and deregulated in each sample separated by category**. Identified metabolites are *m/*z detected either in the sample or in the control. The percentages were calculated by dividing the number of formulas assigned or deregulated in the sample in each category by the number of formulas identified in that sample and multiplying by 100.

**Table S5. Parameters used for bioinformatic analyses.**

**Table S6. Sequencing results and alignment summary**.

